# AKT: Ancestry and Kinship Toolkit

**DOI:** 10.1101/047829

**Authors:** Rudy Arthur, Ole Schulz-Trieglaff, Anthony J. Cox, Jared Michael O’Connell

## Abstract

Ancestry and Kinship Toolkit (AKT) is a statistical genetics tool for analysing large cohorts of whole-genome sequenced samples. It can rapidly detect related samples, characterise sample ancestry, calculate correlation between variants, check Mendel consistency and perform data clustering. AKT brings together the functionality of many state-of-the-art methods, with a focus on speed and a unified interface. We believe it will be an invaluable tool for the curation of large WGS data-sets.

**Availability:** The source code is available at https://illumina.github.io/akt

## 1 Introduction

As whole genome sequencing (WGS) costs decrease, it is becoming common to have re-sequencing data for cohorts of thousands of individuals (Taylor *et al.*, 2015; Lek *et al.*, 2015; Gudbjartsson *et al.*, 2015). Such large cohorts will often have cases of sample duplication, cryptic relatedness and heterogeneous ancestry. These data sets require careful curation before further analysis. In the DNA-microarray world, a range of high-quality tools are available to perform principal component analysis (PCA), kinship coefficient calculation and other routine quality control analyses (Chang *et al.*, 2015). While the algorithms implemented in such tools remain relevant, they require custom formats that are not well suited to WGS data. The conversion between the standard WGS format (VCF/BCF) and these custom formats can be time consuming and error prone for end-users. Additionally, the larger number of rare variants and false-positives in WGS data require some care to handle correctly.

In this note we present *AKT*, a software suite designed to perform routine analyses of large re-sequencing data sets. We envision AKT being applied to large multi-sample BCFs to identify related samples, detect sample swaps and ascertain the spectrum of ancestry in a cohort. Our focus is on speed and simplicity with the hope that this toolkit can become a standard part of the bioinformatician’s arsenal when investigating large cohorts of WGS samples. AKT is freely available under the GPLv3 license. It is implemented in C++ using HTSlib (Li *et al.*, 2009) for fast reading of VCF/BCF files and the Eigen matrix library for matrix manipulations (http://eigen.tuxfamily.org).

## 2 Methods

AKT follows the popular bioinformatics convention of combining many sub-functions into a single binary, analyses are run via: *akt subcommand input.bcf*. Many of the algorithms we describe do not require the entire dense set of variants that will be present in a WGS cohort. Indeed, some of the estimators assume variants are in linkage equilibrium. A standard way of achieving this is to thin variants. We provide thinning/pruning functionality, but this involves decompressing an entire BCF which is time consuming. For example, the final release of the 1000 Genomes Project (1000GP) has 84.8 million variants (The 1000 Genomes Project Consortium, 2015). Our preferred approach is to provide AKT with a pre-determined well behaved sparse set of common SNPs, AKT can then use tabix indexing (Li, 2011) which substantially reduces file reading time. We distribute appropriate site-only VCFs with AKT.

*Fast principal component analysis* PCA is a common method to detect and classify ancestry (Patterson *et al.*, 2006). Plotting the first few principal components will identify large population structure present in a cohort. Reducing a large genotype matrix with *M* markers and *N* samples to principal components requires calculating its singular value decomposition (SVD). Exact SVD is quite slow, *O*(*MN*^2^) when *M>N*. However it is often sufficient to compute an inexact SVD and obtain the first and most important principal components. We implement the very fast randomised approximate SVD routine described in Halko *et al.* (2011). We also provide options to compute the exact SVD using the Jacobi algorithm and to project samples onto pre-computed principal components.

*Kinship coefficients and average IBD sharing* Estimating the proportion of the genome that is identical-by-descent (IBD) between two samples allows us to ascertain the degree of relatedness between them or to check if the samples are duplicates. We calculate the same IBD estimators used in PLINK by default which require population allele frequencies. Users can either estimate frequencies from their data or provide pre-computed frequencies from a reference panel such as 1000GP. The latter option can be especially useful when sample sizes are small. We also provide the option to calculate the KING estimator (Manichaikul *et al.*, 2010) which is robust to population structure and the genetic-relatedness metric which is popular in the mixed effect model community (Yang *et al.*, 2011).

*Detecting cryptic pedigrees* We implement a similar routine to Staples *et al.* (2014). First order relationships (parent-child and sibling) have IBD patterns which allow easy classification. When both parents in a nuclear family are assayed, pedigrees can be reconstructed unambiguously. In cases where only one parent is assayed, a parent-child relationship can be established but not which sample is the parent. Grandparent-grandchild relationships and sibling relationships (when no parents are assayed) can also be detected.

*Other functions* We also include code for data clustering using k++-means, Gaussian mixtures and density based methods (Rodriguez and Laio, 2014), calculation of LD metrics including correlation and LD score (Bulik-Sullivan *et al.*, 2015), transforming principal component projections to ancestry fractions (Zheng and Weir, 2016), tag SNP selection, LD-pruning, simple association testing and profiling of Mendelian inheritance patterns for pedigrees.

## 3 Results

We demonstrate the speed and ease-of-use of AKT on publicly available 1000GP data. We test AKT on two data-sets:

- 1000GP phase 3 release (2504 unrelated samples, 84.8M variants)
- 433 high-coverage samples (including 129 trios and 9 duos, 34.4M variants).

The first data set is perhaps the most commonly analysed WGS cohort, the latter allows us to evaluate the pedigree analysis components of AKT.

Table 1 lists the commands used and respective timings. On the larger *N* = 2504 dataset, PCA took 47 seconds and kinship coefficient calculation took 51 seconds. Re-constructing pedigrees on the smaller data set, took *<* 7 seconds. All trios were correctly reconstructed and all parent-child relationships were identified (we acknowledge this is a fairly straightforward example). Profiling Mendel error rates on these pedigrees across all 34.4M sites took 238 seconds. All results were performed using a single-threaded process (some subcommands accommodate multithreading for faster processing).

**Table 1.** Table 1. Timing results for a subset of AKT functionality on an Intel Xeon E5-2670 CPU. Analysis was performed on the 1000 Genomes Phase 3 BCF (n2504.bcf) and on a separate set of 433 high-coverage samples (n433.bcf). Where appropriate, we perform analysis using a thinned list of 17535 common SNPs (snps.vcf.gz).

| Algorithm | Command line | Time (s) |
| --- | --- | --- |
| PCA | akt pca -R snps.vcf.gz n2504.bcf > n2504.pca | 46.56 |
| kinship | akt kin -R snps.vcf.gz n2504.bcf > n2504.kin | 51.25 |
| Finding cryptic pedigrees | akt kin -R snps.vcf.gz n433.bcf > n433.kin<br>akt relatives n433.kin -p n433.fam | 5.49<br><1 |
| Mendel profile | akt mendel n433.bcf -p n433.fam | 237.65 |

Supplementary section 1 includes some cursory timing comparisons with other popular routines, demonstrating AKT contains competitive implementations. Supplementary section 2 compares our approximate PCA routine to the exact routine available in PLINK and shows the first nine principal components are essentially identical.

## 4 Conclusion

AKT is a convenient tool for for bioinformaticians who routinely deal with large numbers of WGS samples. AKT will help in cases where meta-data about the samples may be missing or unreliable, allowing easy inference of ancestry and relatedness from the data itself. We expect to expand the functionality of AKT with time, however the software will already enable rapid and accurate curation of WGS data. This short analysis gives a feel for the power and speed of AKT for some common problems.

*Funding:* All authors are employees of Illumina Inc., a public company that develops and markets systems for genetic analysis, and receive shares as part of their compensation.

